# Distinct fastigial output channels and their impact on temporal lobe seizures

**DOI:** 10.1101/2021.08.18.456836

**Authors:** Martha L. Streng, Madison Tetzlaff, Esther Krook-Magnuson

**Author notes:** Correspondence: Martha Streng, Ph.D., Department of Neuroscience, University of Minnesota, Jackson Hall, room 6-145, 321 Church Street SE, Minneapolis, MN 55455.

## Abstract

Despite being canonically considered a motor control structure, the cerebellum is increasingly recognized for important roles in processes beyond this traditional framework, including seizure suppression. Excitatory fastigial neurons project to a large number of downstream targets, and it is unclear if this broad targeting underlies seizure suppression, or if a specific output may be sufficient. To address this question, we used the intrahippocampal kainic acid mouse model of temporal lobe epilepsy, male and female animals, and a dual-virus approach to selectively label and manipulate fastigial outputs. We examined fastigial neurons projecting to the superior colliculus, medullary reticular formation, and central lateral nucleus of the thalamus, and found that these comprise largely non-overlapping populations of neurons which send collaterals to unique sets of additional thalamic and brainstem regions, creating distinct, somewhat overlapping, “output channels”. We found that neither optogenetic stimulation of superior colliculus nor reticular formation output channels attenuated hippocampal seizures. In contrast, on-demand stimulation of fastigial neurons targeting the central lateral nucleus robustly inhibited seizures. Our results indicate that fastigial control of hippocampal seizures does not require simultaneous modulation of many fastigial output channels. Rather, selective modulation of the fastigial output channel to the central lateral thalamus, specifically, is sufficient for seizure control. This may provide a means for more selective therapeutic interventions, which provide seizure control while minimizing unwanted side effects. More broadly, our data highlight the concept of specific cerebellar output channels, whereby discrete cerebellar nucleus neurons project to specific aggregates of downstream targets, with distinct functional outcomes.

**Significance statement:** The cerebellum has an emerging relationship with non-motor systems and may represent a powerful target for therapeutic intervention in temporal lobe epilepsy. We find that fastigial neurons project to numerous brain regions via largely segregated output channels, and that excitation of fastigial neurons projecting to the central lateral nucleus of the thalamus, but not the superior colliculus or reticular formation, is sufficient to attenuate hippocampal seizures. These findings illustrate an important conceptual framework: fastigial neurons project to aggregates of downstream targets via distinct output channels, which cannot be predicted simply by somatic location within the nucleus, and these channels have distinct functional outcomes. This nuanced appreciation of fastigial outputs may provide a path for therapeutic interventions with minimized side effects.

## INTRODUCTION

The cerebellum contains more than half of all total neurons in the central nervous system (Andersen et al., 1992), accounts for as much as 20% of the total oxygen consumption of the brain (Howarth et al., 2010), and is reciprocally connected with a large number of cortical and subcortical regions (Ramnani, 2006; Strick et al., 2009; Salmi et al., 2010; Ramnani, 2012; Kipping et al., 2013). Though canonically considered a motor control structure, mounting evidence indicates that the cerebellum is heavily involved in functions beyond this traditional framework (Schmahmann, 1996; Hilber et al., 1998; Leggio et al., 1999; Colombel et al., 2004; Popa et al., 2014; Yu and Krook-Magnuson, 2015; Schmahmann, 2019; Shipman and Green, 2019). Recent work has shown that the cerebellum can profoundly influence hippocampal function, with the ability to modulate hippocampal neuronal dynamics (Choe et al., 2018; Zeidler et al., 2020) and alter hippocampal-dependent behavior (Rochefort et al., 2011; Lefort et al., 2019; Zeidler et al., 2020). The influence of cerebellar dynamics on hippocampal function, in addition to being a topic of great scientific interest, has the potential to meet an urgent translational need in temporal lobe epilepsy (TLE), a disorder characterized by chronic, spontaneous seizures typically arising in the hippocampal formation. TLE is the most common form of epilepsy in adults, but current treatment options have limited efficacy and carry the potential for problematic side effects, leaving 30-40% of epilepsy patients with uncontrolled seizures (England et al., 2012). We recently demonstrated that even very brief optogenetic interventions delivered to excitatory neurons in the cerebellar fastigial nucleus was highly effective at terminating hippocampal seizures in a mouse model of TLE (Streng and Krook-Magnuson, 2020b).

While recent work indicates a lack of a direct, mono-synaptic, connections between the cerebellum and the hippocampus, at least in rodents (Rochefort et al., 2013; Bohne et al., 2019; Krook-Magnuson, 2020), the fastigial nucleus does project to over 60 downstream targets (Fujita et al., 2020). This large number of output targets raises the possibility that seizure suppression via fastigial excitation requires the coordinated modulation of many areas (Eelkman Rooda et al., 2021). Alternatively, a specific output may be sufficient to inhibit seizures, and previous work suggests at least some degree of segregation of cerebellar outputs (Noda et al., 1990; Fuchs et al., 1993; Teune et al., 2000; Zhang et al., 2016; Fujita et al., 2020). Fastigial targets of potential interest for hippocampal seizure control include the thalamus, superior colliculus, and the reticular formation (Angaut and Bowsher, 1970; Batton et al., 1977; Bentivoglio and Kuypers, 1982; Andrezik et al., 1984; Angaut et al., 1985).

The cerebellum has numerous connections with thalamocortical networks (Middleton and Strick, 1998), and the fastigial nucleus in particular projects to several distinct thalamic nuclei (Haroian et al., 1981; Fujita et al., 2020). Deep brain stimulation (DBS) trials targeting the thalamus (albeit the anterior nucleus) have reduced seizures for some patients (Salanova et al., 2015) and activation of neurons in the deep cerebellar nuclei disrupt spike-and-wave discharges observed during thalamocortical absence seizures (Kros et al., 2015; Eelkman Rooda et al., 2021). Cerebellar connections to intralaminar and midline thalamic nuclei may be especially relevant for TLE, as they have been implicated in regulating limbic seizures (Wicker and Forcelli, 2016; Feng et al., 2017). While much of the current seizure literature has focused on the central lateral (CL) thalamus in the context of consciousness (Gummadavelli et al., 2015; Kundishora et al., 2017; Xu et al., 2020), the CL is of particular interest to us in the context of cerebellar mediated seizure suppression: its activity is depressed (and bursty) during focal limbic seizures (Feng et al., 2017), and it provides a potential route (via the anterior cingulate) from the cerebellum to the hippocampus (Van der Werf et al., 2002; Rajasethupathy et al., 2015).

Another candidate region of interest is the superior colliculus (SC), which in addition to receiving fastigial input (Roldan and Reinoso-Suarez, 1981; Fujita et al., 2020) is proposed to play a role in regulating seizure activity (Garant and Gale, 1987; Dean and Gale, 1989; Weng and Rosenberg, 1992), and can inhibit seizures in multiple animal models (Gale et al., 1993; Soper et al., 2016). The fastigial nucleus also has extensive projections to the reticular formation (Andrezik et al., 1984; Zhang et al., 2016; Fujita et al., 2020), a collection of nuclei important for controlling brain states (Moruzzi and Magoun, 1949; Jones, 2003) which could potentially underlie seizure control (Ewell et al., 2015; Khan et al., 2018; Purnell et al., 2018; Streng and Krook-Magnuson, 2020a). Clearly, there are several strong candidate regions for mediating the seizure inhibition seen with on-demand fastigial stimulation, if concurrent excitation of multiple downstream targets is not required.

We therefore set out to examine fastigial outputs to the central lateral thalamus, the superior colliculus, and the medullary reticular formation, to determine 1) if these areas are targeted by separate fastigial neurons, 2) if fastigial neurons targeting these areas also target other areas (and if so, which), and 3) if any of these output channels, in isolation, is able to inhibit seizures. Using a dual viral targeting strategy, we labeled populations of fastigial neurons that project to the CL, SC, or reticular formation. We find that these neurons represent largely distinct populations and project to generally segregating additional downstream areas. Using the intrahippocampal kainate mouse model of temporal lobe epilepsy, and on-demand optogenetic manipulation of specific output pathways upon online detection of spontaneous seizures (Armstrong et al., 2013), we further find that cerebellar control over hippocampal seizures can be achieved via activation of a specific output channel, rather than requiring broader network effects. Specifically, excitation of CL-projecting fastigial neurons, but not the other two output pathways examined, is able to robustly attenuate hippocampal seizures. Together, our results illustrate nuanced fastigial nucleus output channels with important, potentially clinically relevant, different functional consequences.

## MATERIALS AND METHODS

### Ethical approval

All experimental protocols were approved by the University of Minnesota’s Institutional Animal Care and Use Committee.

### Animals

For all experiments, mice were bred in-house and had *ad libitum* access to food and water in all housing conditions. Black-6 mice (C57BL/6J; Jackson Laboratory stock 000664) were used for initial examination of broad fastigial projections, for output channel specific labeling and modulation, and for a subset of terminal stimulation experiments. Mice expressing Cre selectively in VGluT2-expressing neurons (Vong et al., 2011) (B6J.129S6(FVB)-Slc17a6^tm2(cre)Lowl^/MwarJ; Jackson Laboratory stock 028863) were also utilized for on-demand terminal stimulation experiments.

Animals were sexed at the time of weaning on the basis of external genitalia. Both male and female mice were used for all experiments. While experiments were not powered to test for sex differences, no trends of sex differences were observed. Until optical fiber and electrode implantation, animals were housed in standard group housing conditions in the Research Animal Resources animal facility at the University of Minnesota. Following implantation, animals were singly housed, and experiments were performed while housed in investigator managed housing. In all conditions, animals were allowed ad libitum access to food and water, and were on a 12 hour light; 12 hour dark (/low red light) cycle.

### Stereotactic surgeries

#### Viral targeting

For all experiments, AAV serotype 9 was used for opsin expression in fastigial neurons due to its optimal expression in the fastigial nucleus with no apparent retrograde expression (Streng and Krook-Magnuson, 2020b). All injections were performed in adult mice (postnatal day 45 or later).

For initial characterization of fastigial fibers, Black-6 mice were injected with 120nL of virus encoding GFP in a Cre-independent manner (AAV9-CAG-GFP, titer of 2×10e^12^, UNC vector core lot #AV5221, provided to UNC by Edward Boyden) via a Hamilton Neuros syringe into the left cerebellar fastigial nucleus (6.48 posterior, 0.75 left, 3.7 mm ventral from bregma) under isoflurane anesthesia.

For experiments examining specific fastigial output channels, Black-6 mice were first injected with 120nL of a retrograde virus encoding Cre (AAVrg-Ef1a-mCherry-IRES-Cre, titer of 1.37 x 10^13, Addgene viral prep # 55632-AAVrg, provided to Addgene by Karl Deisseroth (Fenno et al., 2014)) in either the central lateral nucleus (CL) (1.34 mm posterior, 0.75 mm right, 3.0 mm ventral from bregma), superior colliculus (SC) (4.0 mm posterior, 1.0 mm right, 2.5 mm ventral from bregma), ventral medullary reticular nucleus (MdV) (7.5 mm posterior, 0.36 mm right, 4.0 mm ventral from bregma), or ventral lateral nucleus (VL) (1.25 mm posterior, 0.96 mm right, 3.5 mm ventral from bregma). Following retrograde viral injection, the contralateral fastigial nucleus (6.48 posterior, 0.75 left, 3.7 mm ventral from bregma) was injected with virus encoding Channelrhodopsin in a cre-dependent manner (AAV9-EF1a-double floxed-hChR2(H134R)-EYFP-WPRE-HGHpA, titer of 2.2×10^13^ Addgene viral prep #202198-AAV9, lot #V22125, provided to Addgene by Karl Deisseroth)(Gradinaru et al., 2007) consistent with our previously published methods for successful viral targeting of this nucleus (Streng and Krook-Magnuson, 2020b). After every injection, the syringe was held in place for a minimum of 10 minutes before being withdrawn. On-demand interventions and/or characterization of fibers were conducted a minimum of 6 weeks post viral injection. Mice with off target expression were excluded from analyses (n=1 animal).

For experiments targeting fastigial terminals, Black-6 mice were injected with virus encoding Channelrhodopsin fused to green fluorescent protein (AAV9-CAG-ChR2-GFP, titer of 2.1×10e^12^, UNC vector core lot #AV5406D, provided to UNC by Edward Boyden) or VGluT2-Cre mice were injected with virus encoding Channelrhodopsin in a Cre-dependent manner (AAV9-EF1a-double floxed-hChR2(H134R)-EYFP-WPRE-HGHpA, titer of 2.2×10^13^, Addgene viral prep #202198-AAV9, lot #V22125, provided to Addgene by Karl Deisseroth)(Gradinaru et al., 2007). For opsin negative control experiments, Black-6 mice were injected with virus encoding GFP alone (AAV9-CAG-GFP, titer 2×10^12^, UNC vector core lot #AV5221, provided by Edward Boyden).

#### Epilepsy induction

Procedures for epilepsy induction using the mouse unilateral intrahippocampal kainic acid model of TLE largely followed previously published protocols (Cavalheiro et al., 1982; Bouilleret et al., 1999; Bragin et al., 1999; Krook-Magnuson et al., 2013; Krook-Magnuson et al., 2014; Streng and Krook-Magnuson, 2020b). A minimum of two weeks post viral injection(s), mice were injected with 100nL of kainic acid (KA) unilaterally into the right dorsal hippocampus (2.0 mm posterior, 1.25 mm right, 1.6 mm ventral from bregma) under isoflurane anesthesia (2%), as done in our previous work (Streng and Krook-Magnuson, 2020b). Animals were removed from isoflurane a maximum of five minutes post injection (Bar-Klein et al., 2016). In this model, spontaneous recurrent electrographic seizures emerge from the damaged hippocampus, providing a strong model of pharmocoresistant (Riban et al., 2002; Klein et al., 2015) temporal lobe epilepsy with hippocampal sclerosis (Bouilleret et al., 1999; Riban et al., 2002; Levesque and Avoli, 2013; Zeidler et al., 2018). Only animals that showed spontaneous hippocampal seizures weeks after KA injection, as determined by video EEG monitoring, were included for on-demand optogenetic interventions.

#### Electrode and fiber implantation

A minimum of 1 week post kainic acid injection, mice were implanted with a twisted wire bipolar (local reference, differential) electrode (PlasticsOne) ipsilateral to the site of kainate (2.6 mm posterior, 1.75 mm right, 1.6 mm ventral from bregma). For fastigial output channel targeting experiments, mice were additionally implanted with optical fibers targeting the left fastigial nucleus (6.48 mm posterior, 0.75 mm left, 2.5 mm ventral from bregma). For fastigial terminal targeting experiments, mice were implanted with optical fibers targeting the right central lateral nucleus of the thalamus (1.34 mm posterior, 0.75 mm right, 2.75 mm ventral from bregma). Implants were secured to the skull using screws and dental cement following previous protocols (Armstrong et al., 2013; Krook-Magnuson et al., 2013) and allowed to recover a minimum of five days prior to seizure monitoring and closed-loop interventions.

#### Post-operative care

For all surgical procedures, post-operative care consisted of recovery from anesthesia on a heating pad with regular visual inspection, followed by daily post-operative monitoring for a minimum of three days to inspect comfort level and healing of the surgical site. Neopredef powder was applied to the closed incisions as a topical antibiotic and analgesic. In the case of kainic acid injection, no additional post-operative analgesics were given. For viral injections and implantations, carprofen was administered subcutaneously (5mg/kg) immediately prior to surgery. For implantations, pre- and post-operative ibuprofen was also administered orally (50-80mg/kg/day in water) as an additional analgesic.

### Closed-loop seizure detection and interventions

A minimum of five days post implantation, animals were placed in investigator managed housing for chronic video and LFP recordings. Detection of electrographic seizures and on-demand optogenetic interventions generally followed previously published protocols (Armstrong et al., 2013; Krook-Magnuson et al., 2013; Streng and Krook-Magnuson, 2020b). Hippocampal LFP was recorded via electrical patch cords through an electrical commutator (PlasticsOne), amplified (Brownlee), digitized (National Instruments), and analyzed in real time by custom MATLAB seizure detection software. A version of this software is available for download through Armstrong *et al*, 2013 (Armstrong et al., 2013). Optogenetic interventions consisting of three seconds of pulsed light delivery (50 msec on, 100 msec off) (Streng and Krook-Magnuson, 2020b) were triggered for 50% of events in a random fashion using this custom closed-loop MATLAB software, and were delivered via LEDs (473 nm) through optical patch cords (Thor labs). Stimulation parameters were selected due to their efficacy in robustly attenuating hippocampal seizures when targeting the fastigial nucleus more broadly (Streng and Krook-Magnuson, 2020b) or the cerebellar cortex (Krook-Magnuson et al., 2014), and were originally selected to minimize movement side-effects (Krook-Magnuson et al., 2014). Post-hoc measurements of light delivery indicated an average LED power of 1.5 +/− 0.6mW at the tip of the implanted optical fiber.

### Tissue harvesting and imaging of fibers

In order to characterize viral expression and confirm appropriate optical fiber targeting, after completion of on-demand interventions, mice were either deeply anesthetized with 5% isoflurane and decapitated or perfused with 0.1M phosphate buffer followed by 4% paraformaldehyde. Brains were subsequently harvested and fixed in 4% paraformaldehyde. Sagittal brain sections of 50µm were collected in 0.1M phosphate buffer using a vibratome (Leica VT1000S). After sectioning, every third section was mounted with Vectashield mounting media with DAPI and covered with glass coverslips. Sagittal sections were initially visualized with epifluorescence microscopy (Leica DM2500) to identify sections with structures of interest for confocal imaging. The approximate coordinates of the sagittal slices and structures of interest were determined using the Paxinos mouse brain atlas, third edition (Franklin and Paxinos, 2007) in combination with DAPI and brightfield microscopy, which allowed for the discernment of areas of interest, including thalamic nuclei. Confocal imaging was performed on an Olympus FluoView FV1000 BX2 upright confocal microscope (University Imaging Center, University of Minnesota), with fields of view first selected using the DAPI signal to determine location (that is, blinded to the specific eYFP fiber locations/densities). Medial lateral coordinates for the structures of interest are as follows: SC (0.72 mm right), MdV (0.48 mm right), CL (0.72 mm right), medial dorsal nucleus, lateral part (MDL; 0.72 mm right), medial dorsal nucleus, central part (MDC; 0.48 mm right), ventral medial nucleus (VM, 0.72 mm right), ventral lateral nucleus (VL, 0.72 mm right), parafascicular nucleus (PaF, 0.6 mm right), zona incerta (ZI, 1.44 mm right), mesencephalic reticular formation (mRT, 1.56 mm right), laterodorsal tegmental nucleus (LDTg, 0.48 mm right), lateral periaqueductal grey (lPAG, 0.84 mm right), ventral lateral periaqueductal grey (vlPAG, 0.48 mm right), nucleus reticularis pontis caudalis (PnC, 0.48 mm right), parvocellular reticular nucleus (PCRT, 1.20 mm right), and spinal vestibular nucleus (SpVe, 1.44 mm right). Representative images were adjusted for brightness and contrast.

### Quantification of fibers

After confocal images of eYFP labeled fibers in structures of interest were taken from FN-SC, FN-MdV, and FN-CL mice (one image taken per location from 50µm sagittal sections from three animals from each group), neurites within the imaged region of interest were measured using Fiji’s simple Neurite Tracer by tracing each neurite and the corresponding branches as separate paths (note that quantification of fibers was done for only a subset of areas showing expression). For the creation of a simplified summary schematic, the relative magnitude of fastigial fibers in a given location was assessed by comparing to the maximum observed in that area (for this, we additionally quantified fibers in a mouse injected with virus for broad expression of ChR2-GFP). Projections from a particular FN output channel to a region of interest were included in the summary schematic if the average total path length was at least 15% of maximum and was present at a minimum 15% level in at least two of the three animals quantified, with line thickness proportional to the percent of maximum of the average total path length for that output location.

### Immunohistochemistry

Our procedures for VGluT2 immunohistochemistry largely follow our previously published protocol (Streng and Krook-Magnuson, 2020b). Briefly, every third sagittal section containing the downstream target of interest was blocked with 10% bovine serum and 0.5% triton diluted in TBS, followed by overnight incubation with primary antibody for VGluT2 at 4°C (Millipore Sigma, 1:1000 diluted in TBS containing 2% bovine serum and 0.4% triton). Tissue was then rinsed and incubated for 2 hours with Alexa Fluor 594 anti-guinea pig (Jackson, 1:500 diluted in TBS containing 2% bovine serum and 0.4% triton). Following secondary incubation, tissue was mounted with Vectashield mounting media with DAPI and visualized with epifluorescence microscopy (Leica DM2500) and confocal microscopy (Olympus FluoView FV1000 BX2) to confirm colocalization with eYFP+ fibers.

For osteopontin (SPP1) immunohistochemistry, every third sagittal section of tissue containing the left fastigial nucleus was blocked with 5% bovine serum albumin for one hour at room temperature prior to incubation in primary antibody goat anti-osteopontin overnight at 4°C (1:300, R&D Systems AF808; RRID:AB_2194992). Tissue was then rinsed and incubated in donkey anti-goat Alexa Fluor 594 for 2 hours at room temperature (1:500, ThermoFisher Scientific A-11058). Following secondary incubation, tissue was mounted with Vectashield mounting media with DAPI and viewed for colocalization of virally labeled and immunolabeled cell bodies using epifluorescence microscopy (Leica DM2500). Additional sections from an additional animal (without immunohistochemistry) were also utilized for additional mapping of virally-labeled somatic locations.

### Statistical analyses

Seizure duration after the time of trigger and time to next seizure were analyzed off-line using a combination of manual and automated methods consistent with our previous methods (Streng and Krook-Magnuson, 2020b). Software for automated analysis is available for download through github (https://github.com/KM-Lab/Electrographic-Seizure-Analyzer) (Zeidler et al., 2018). A minimum of 100 seizure events per animal per condition were processed automatically based on user-identified characteristics of spikes including amplitude, peak width, spike frequency and deflection (positive, negative, or both), and the resulting post-detection seizure duration of all events were confirmed via manual inspection. Post-detection seizure duration distributions between light and no light conditions were compared in each animal using two-sample Kolmogorov-Smirnov tests. To assess effectiveness of intervention at the group level, two approaches were taken. First, a summative histogram of events was created, using an equal number of events per animal, and the resulting distributions of light and no-light event durations compared using two-sample Kolmogorov-Smirnov tests. Second, average post-detection seizure durations (one value per condition per animal) were compared, and for visualization in figures, post-detection seizure durations with light intervention were normalized by dividing by duration without intervention (no light internal controls for the same animal), and expressed as a percent. A *p* value < 0.05 was considered statistically significant. Statistical analyses were conducted using MATLAB. Values are presented as mean ± SEM.

## RESULTS

### Fastigial outputs target numerous downstream structures

On-demand excitation of the fastigial nucleus can robustly inhibit hippocampal seizures (Streng and Krook-Magnuson, 2020b). However, how the fastigial nucleus can produce such a profound influence on seizure activity is unclear. The fastigial projects to numerous downstream regions (Angaut and Bowsher, 1970; Batton et al., 1977; Bentivoglio and Kuypers, 1982; Teune et al., 2000; Fujita et al., 2020). We first visualized fastigial projections to potential downstream targets of interest using a similar viral targeting strategy to our on-demand fastigial excitation work (Streng and Krook-Magnuson, 2020b). Specifically, we injected Black-6 mice with an AAV to achieve expression of the fluorescent protein GFP in neurons of the fastigial nucleus (AAV9-CAG-GFP; Fig. 1A). With this approach, we labeled fastigial neurons in both rostral and caudal portions of the fastigial nucleus (Fig. 1B-C), consistent with our previously published results (Streng and Krook-Magnuson, 2020b).

**Figure 1.**
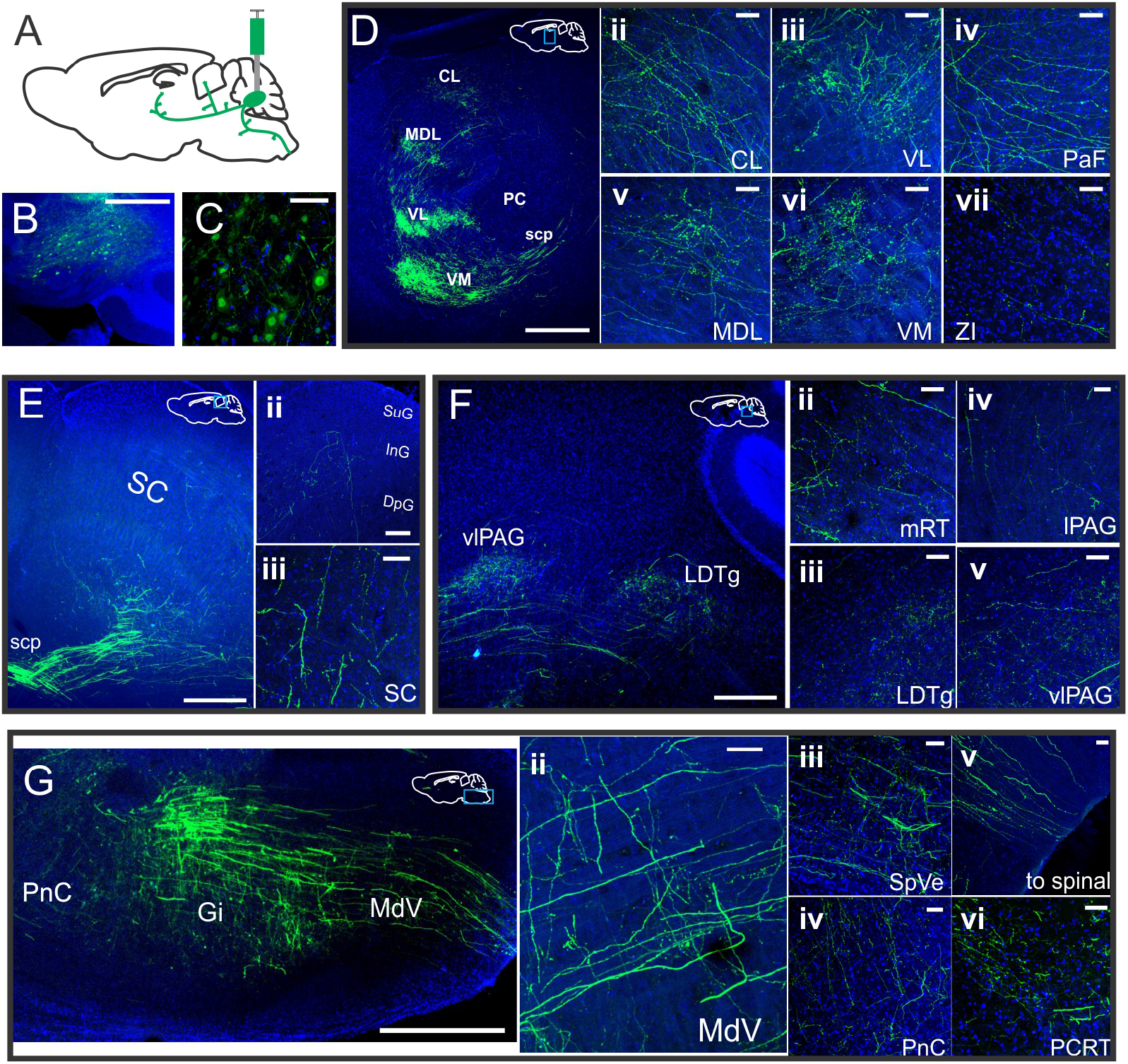
Viral labeling illustrates a large number of downstream targets of fastigial neurons. A) Schematic of injection targeting, in which mice are injected with an AAV to induce the expression of GFP in fastigial neurons for broad labeling of fastigial outputs. B-C) GFP expression in fastigial neurons at two different magnifications. D) Six weeks post injection, GFP+ fastigial fibers are visible in the thalamus (D) including central lateral (ii), ventral lateral (iii), parafascicular (iv), medial dorsal (v), ventral medial (vi) and zona incerta (vii) nuclei. Blue box over white schematic indicates approximate region of focus. E) Fastigial fibers are also visible in the superior colliculus, including both deep and intermediate layers (ii-iii). F) In the midbrain, fastigial fibers are observed in the mesencephalic reticular formation (ii), laterodorsal tegmental nucleus (iii), as well as both the lateral (iv) and ventral lateral (v) periaqueductal gray. G) Extensive fastigial fibers are visible in the brainstem, including the medullary reticular formation (ii), vestibular nuclei including the spinal vestibular nucleus (iii), and pontine reticular nuclei (iv, vi). Fibers are also visible in more caudal regions of the brainstem, likely travelling to the spinal cord (v). CL: central lateral nucleus, MDL: medial dorsal nucleus, lateral part, VL: ventral lateral nucleus, VM: ventral medial nucleus, PC: paracentral nucleus, SC: superior colliculus, scp: superior cerebellar peduncle, PaF: parafascicular nucleus, ZI: zona incerta, SC: superior colliculus, SuG: superior gray layer, InG: intermediate gray layer, DpG: deep gray layer, vlPAG: ventral lateral periaqueductal gray, LDTg: laterodorsal tegmental nucleus, mRT: mesencephalic reticular nucleus, lPAG: lateral periaqueductal gray, PnC: nucleus reticularis pontis caudalis, Gi: gigantocellular reticular nucleus, MdV: medullary reticular nucleus (ventral part), SpVe: spinal vestibular nucleus, PCRT: parvocellular reticular nucleus. All images are taken from sagittal sections; see methods for approximate medial lateral coordinates for each region. Scale bars: 200 µm (B); 50µm (C); 500 µm (D, G); 250 µm (E, Eii, F); 50µm (Dii-vii, Eiii, Fii-v, Gii-vi)

Matching previous reports (Haroian et al., 1981; Asanuma et al., 1983; Angaut et al., 1985; Gornati et al., 2018; Fujita et al., 2020), we found widespread axonal projection targets in numerous regions (Fig. 1), including three areas of particular interest regarding potential seizure suppression: the central lateral nucleus of the thalamus (CL, Fig 1Dii), superior colliculus (SC, Fig. 1E), and medullary reticular formation (MdV, Fig 1Gii). As expected, fastigial terminals were visible in multiple other thalamic nuclei (Fig 1D), including the ventral lateral (Fig. 1Diii), parafascicular (Fig. 1Div), medial dorsal (Fig. 1Dv), ventral medial (Fig. 1Dvi), and zona incerta nuclei (Fig. 1Dvii). In the midbrain, in addition to the superior colliculus (Fig. 1E), expression was also observed in other mibrain nuclei (Fig. 1F) including the mesencephalic reticular formation (Fig. 1Fii), laterodorsal tegmental nucleus (Fig. 1Fiii), and both lateral and ventral lateral portions of the periaqueductal grey (Fig. 1Fiv-v). In the brainstem, fibers were observed in multiple regions of the reticular formation (Fig. 1G, Gii, iv-vi) and vestibular nuclei (Fig. 1Giii). We also observed fastigial fibers traveling caudally through the medulla, likely comprising cerebello-spinal tracts (Fig. 1Gv). Only a subset of regions with GFP+ terminals are presented here; we observed a large number of projection targets, consistent with a recent report identifying over 60 downstream targets (Fujita et al., 2020). The expression pattern after broad targeting of the fastigial nucleus gave us a reference point for our next experiments looking at specific outputs using a dual virus approach.

### Fastigial outputs to the superior colliculus, medullary reticular formation, and central lateral nucleus comprise distinct populations

To examine if neurons projecting to our three areas of interest (SC, MdV, and CL thalamus) represented distinct neuronal populations, we utilized a dual viral strategy to achieve expression of ChR2-eYFP in populations of fastigial neurons defined by their projection targets. Specifically, mice were injected first in the downstream target of interest with an AAV designed to provide retrograde expression of Cre (retroAAV-Cre) (Tervo et al., 2016), followed by injection of the contralateral fastigial nucleus with a Cre-dependent virus for ChR2-eYFP expression (Gradinaru et al., 2007). The result of these dual injections is ChR2-eYFP expression only in fastigial neurons projecting to the downstream target of interest (Fig. 2A, B, C). We injected retroAAV-Cre into either the SC (Fig. 2Aii), MdV (Fig. 2Bii), or CL thalamus (Fig. 2Cii), resulting in labeling of fastigial neurons that project to the SC, MdV, or CL, henceforth referred to as FN-SC, FN-MdV, and FN-CL neurons, respectively.

**Figure 2.**
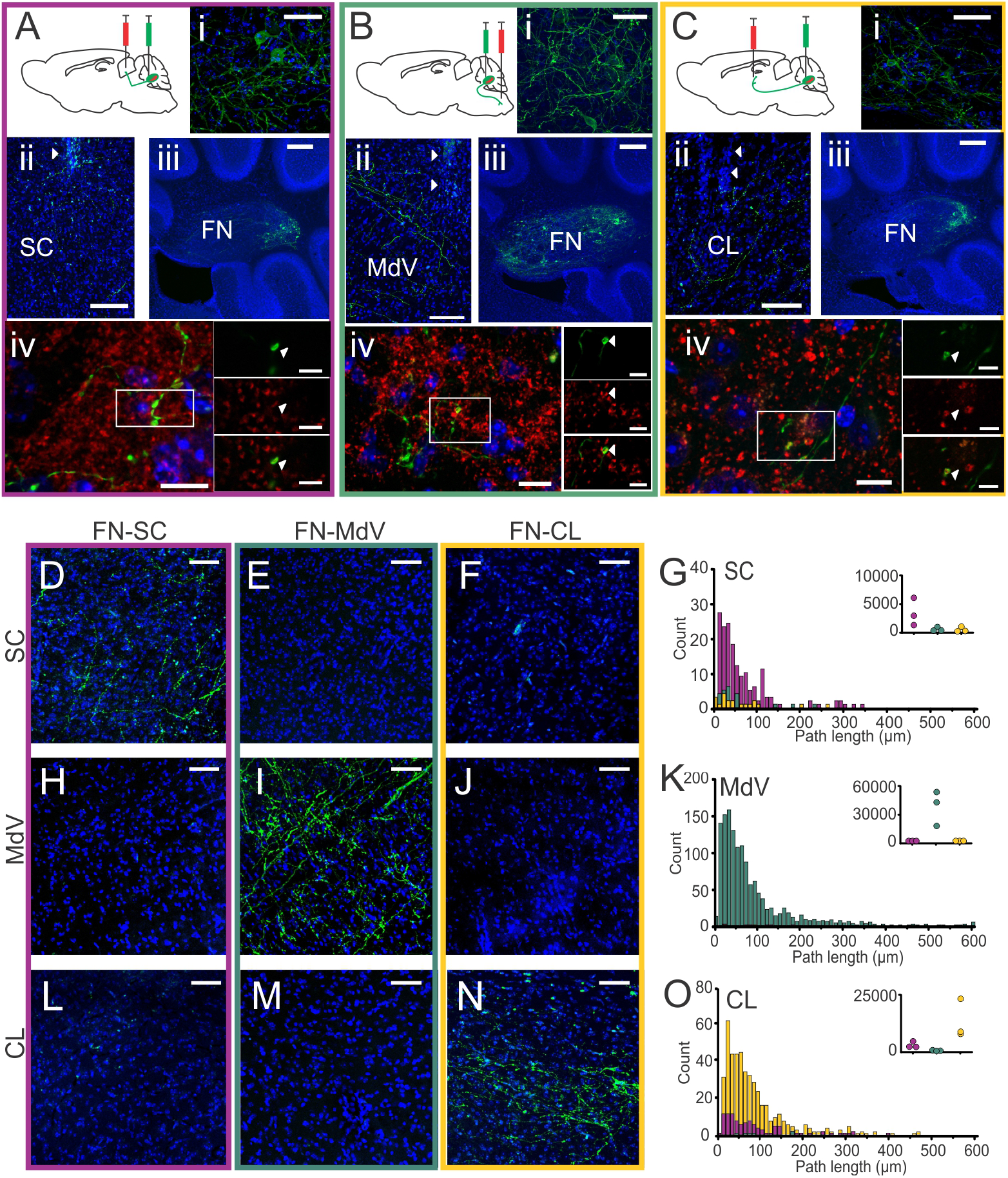
A dual-virus targeting strategy reveals fastigial output channels to segregated projection targets. A) Schematic of injection targeting for FN-SC neurons. Mice are injected with a virus for retrograde expression of Cre (red syringe) in the superior colliculus followed by injection of a virus for Cre-dependent expression of ChR-eYFP in the fastigial nucleus (green syringe). Dual injection results in labeling of fastigial neurons (i) after injection to superior colliculus (ii, arrows denote injection tract). FN-SC cell bodies expressing ChR-eYFP are observed in the caudal fastigial nucleus (iii), and VGluT2+ (shown in red) fastigial fibers (green) are present in the superior colliculus (iv). B) same as for (A), but for MdV injection targeting, in which fastigial cell bodies (i) are labeled after retro-virus injection in the MdV (ii). FN-MdV cell bodies expressing ChR-eYFP are observed in both rostral and caudal portions of the fastigial nucleus (iii), and VGluT2+ fastigial fibers are present in the MdV (iv). C) Same as for (A-B), but for CL retro-virus injection targeting. D-F) Extensive fastigial fibers are observed in the SC for FN-SC, but not FN-MdV nor FN-CL neurons. G) Histogram of path lengths in the SC for FN-SC (magenta), FN-MdV (teal), and FN-CL (yellow) neurons (n = 3 animals per group, inset indicates total path lengths for each individual animal). H-J) Extensive fastigial fibers are observed in the MdV for FN-MdV, but not for FN-SC or FN-CL, neurons. K) Histogram of path lengths and inset dot plots of total path length for each individual animal in the MdV for FN-SC, FN-MdV, and FN-CL neurons. L-N) Dense fibers are observed in the CL for FN-CL, but not SC or MdV, neurons. O) Histogram of path lengths and inset dot plots of total path length for each individual animal in the CL for FN-SC, FN-MdV and FN-CL neurons. All images are taken from sagittal sections. Scale bars: 20µm (Ai, Bi, Ci); 100µm (Aii, Bii, Cii); 200µm (Aiii, Biii, Ciii); 10µm (Aiv, Biv, Civ, insets 2µm) 50µm (D-F, H-J, L-N).

FN-SC labeled neurons (Fig. 2Ai) were located in the caudal portion of the fastigial nucleus (Fig. 2Aiii, compare to Fig. 1B; Fig. 3D). As expected, FN-SC neurons had strong labeling of terminals in the SC (Fig. 2Aiv, D,G). Importantly, fibers from FN-SC neurons were largely absent from the MdV (Fig. 2H,K) and CL (Fig. 2L,O). Conversely, FN-MdV neurons (Fig. 2Bi) were located throughout rostral and caudal portions of the fastigial nucleus (Fig. 2Biii, Fig. 3E), with some labeling extending into the interposed nucleus (Fig. 3E). Fibers from FN-MdV neurons were strongly present in the MdV (Fig. 2Biv, I, K), but not SC (Fig. 2E, G) or CL (Fig. 2M, O). Similar to FN-SC neurons, FN-CL neurons (Fig 2Ci) were preferentially located in the caudal portion of the fastigial nucleus (Fig. 2Ciii, Fig. 3F). Similar independence of expression was observed for FN-CL, with heavy labeling of terminals present in the CL (Fig. 2Civ, N-O), but not SC (Fig. 2F-G) or MdV (Fig. 2J-K). This suggests that FN-SC, FN-MdV, and FN-CL cells represent distinct populations of fastigial neurons. Note also that as all three populations included at least some caudal labelling, full separation of these populations based solely on somatic location would not be possible.

**Figure 3.**
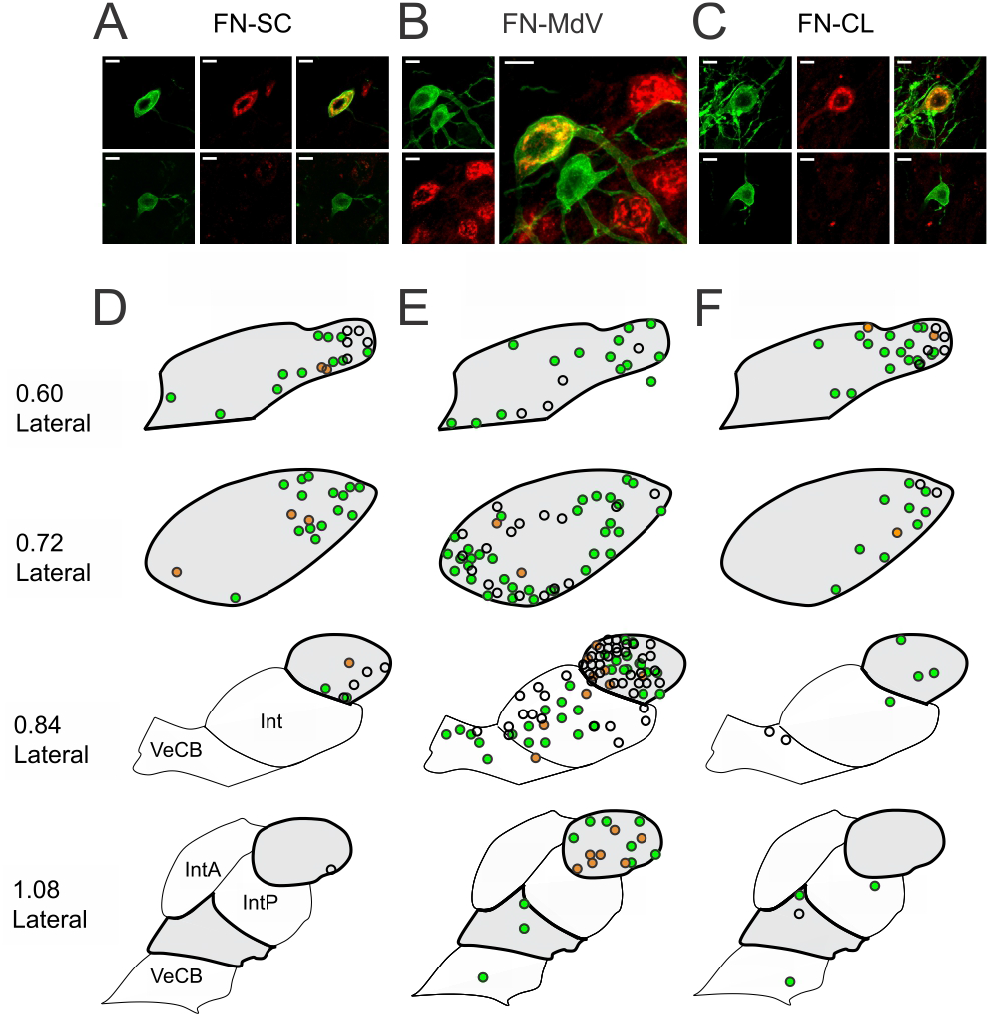
Anatomical characterization of FN-SC, FN-MdV, and FN-CL neurons. A-C) Osteopontin (SPP1) is not a reliable marker of FN-SC, FN-MdV, nor FN-CL neurons. A) Example SPP1-immunopositive (top row; SPP1-immunolabeling in red, eYFP-ChR2 expression in green; right: colocalization) and SPP1-immunonegative (bottom row) FN-SC neuron. B) Example SPP1+ and SPP1- FN-MdV neurons. Note that SPP1+/eYFP+ (yellow in overlay), SPP1+/eYFP- (red in overlay), and SPP1-/eYFP+ (green in overlay) neurons are visible in the same location (field of view) within the FN. C) Example SPP1+ (top row) and SPP1- (bottom row) FN-CL neurons. D-F) FN-SC neurons (D) and FN-CL neurons (F) are primarily located in the caudal FN, whereas FN-MdV neurons (E) are found in both rostral and caudal portions. FN denoted by thick outline and gray shading. Circles denote eYFP labeled cells (n = 3 animals total). For tissue assessed for SPP1 (n = 2 animals each), eYFP cells displaying SPP1+ immunolabeling are filled orange, and eYFP cells negative for SPP1-immunolabeling are filled green. Scale bars: 10µm. Int: interposed nucleus, IntA: Interposed nucleus, anterior part, IntP: interposed nucleus, posterior part, VeCB: vestibulocerebellar nucleus.

As an additional step, we characterized whether FN-SC, FN-MdV, or FN-CL neurons expressed the molecular marker osteopontin (SPP1), which was recently characterized as a potential marker for subpopulations of fastigial neurons (Fujita et al., 2020). We found both SPP1-immunopositive (SPP1^+^) and SPP1-immunonegative (SPP1^−^) fastigial neurons in all three populations (Fig. 3A-C). However, the majority of neurons were SPP1-immunonegative, with only 19.6% (9/46 cells), 20.2% (19/143 cells), and 7.3% (3/41 cells) immunopositive for SPP1 for FN-SC, FN-MDV, and FN-CL neurons, respectively (Fig. 3D-F). While a smaller proportion of FN-CL cells were co-labeled, compared to FN-SC or FN-MdV neurons, this was not statistically significant (p=0.14 and p=0.07, respectively; Pearson’s chi square). These results suggest that SPP1 is not a reliable marker for FN-SC, FN-MDV, or FN-CL neurons.

Importantly, the location of fibers indicates that our dual labeling approach effectively labels three distinct populations of fastigial neurons, in which fastigial neurons that project to the CL nucleus do not project to the SC or MdV, and so on. This is predominantly in keeping with recently reported findings (Fujita et al., 2020), and provided us with two important experimental benefits. First, given that distinct populations of neurons were labeled, this approach provided a means to directly target FN-SC, FN-MdV, or FN-CL somata for circuit dissection of seizure suppression, avoiding caveats associated with strategies optogenetically targeting terminals (e.g., insufficient light coverage). Additionally, this labeling approach allowed us to examine which (if any) other downstream regions each of these populations project to, providing greater insight into the circuit design of fastigial outputs.

We found that fastigial neurons that projected to the SC, MdV, or CL did, indeed, also project to other areas, and that for many of these areas, segregation of these three output channels was largely maintained (Figures 4-7).

**Figure 4.**
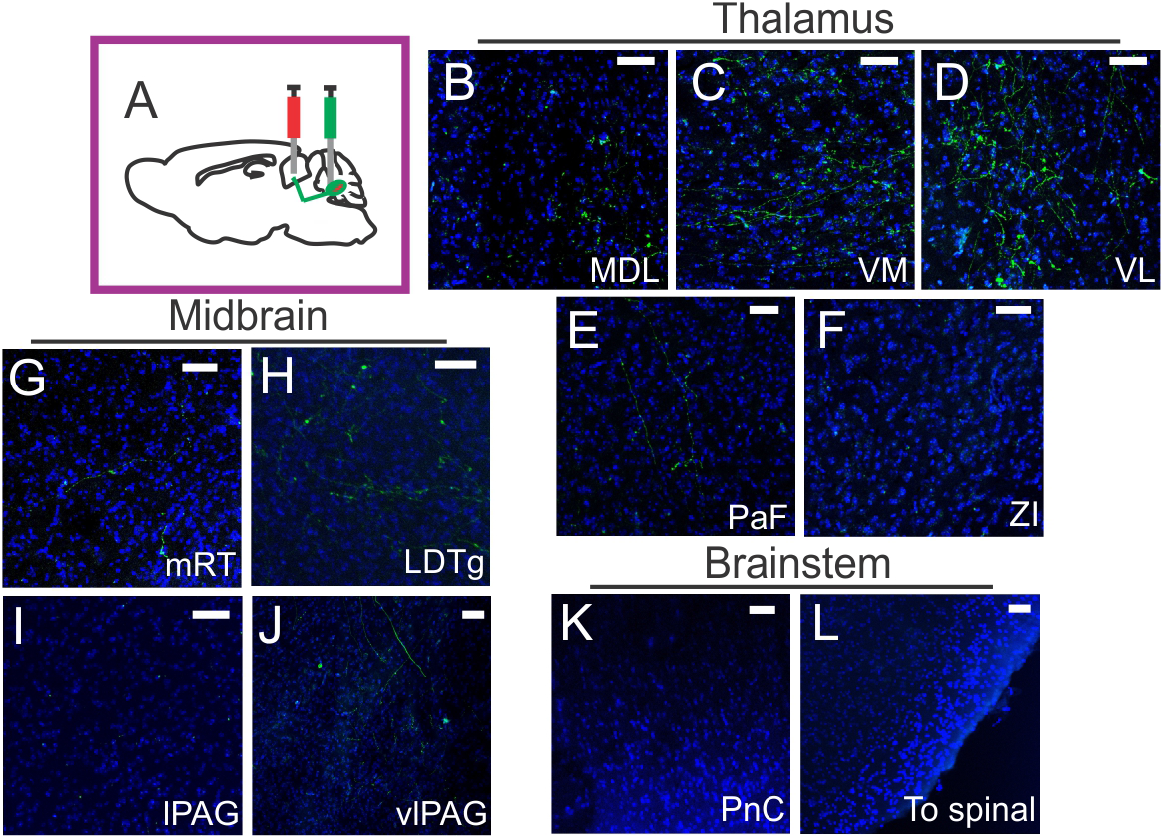
FN-SC neurons send collaterals to distinct thalamic and midbrain nuclei. A) Schematic of injection targeting for FN-SC neurons. In addition to strong fibers in the SC (Figure 2D), FN-SC neurons display some fibers in the thalamus (B-F), especially in the VL nucleus (D). F) No fibers are observed in the zona incerta (ZI). In the midbrain, sparse fibers are observed in some regions including the mRT (G), LDTg (H), and vlPAG (J). Little to no fibers are observed in lPAG (I) or the brainstem, including the PnC (K). There is also no evidence of FN-SC neurons sending fibers to the spinal cord (L). All images are taken from sagittal sections. Abbreviations are as for Fig. 1. Scale bars 50µm.

**Figure 5.**
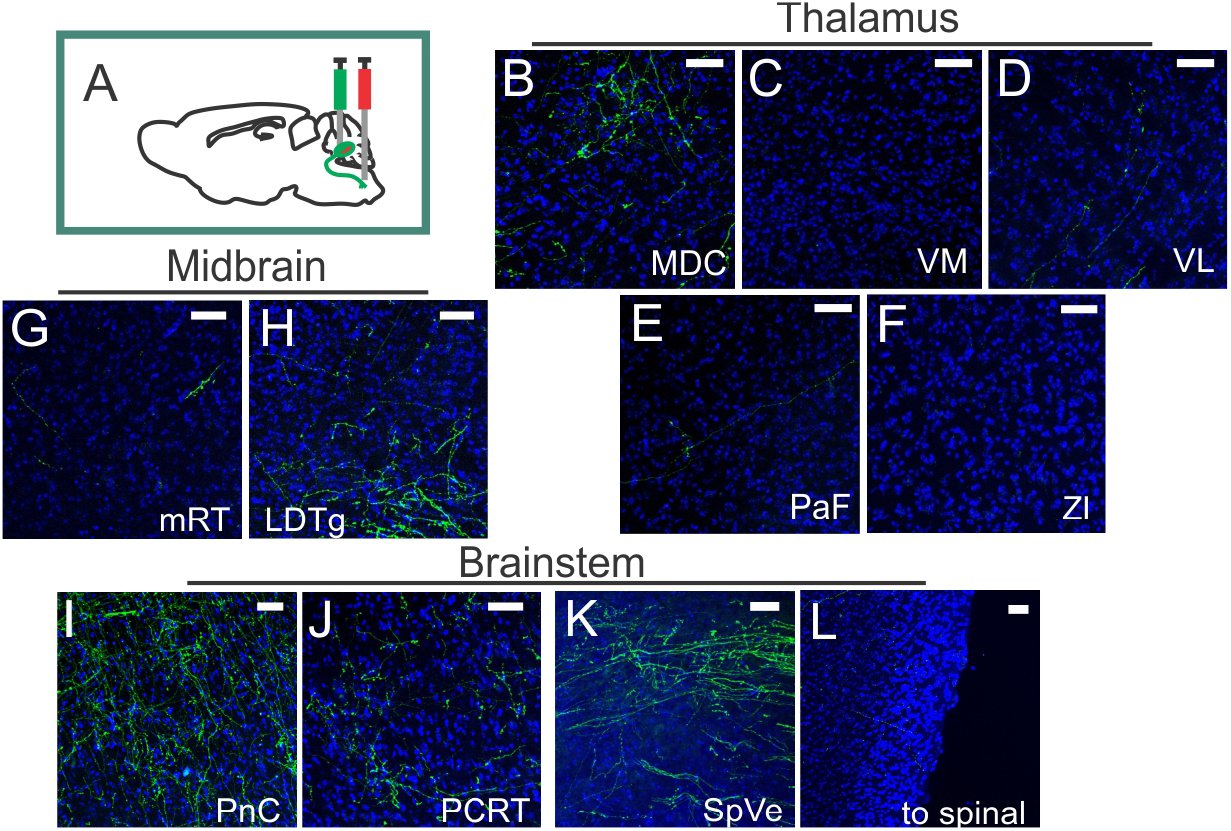
Collaterals from FN-MDV neurons have strong projections to several areas, but are largely restricted to the midbrain and brainstem. A) Schematic of injection targeting for FN-MdV neurons. B-F) FN-MdV fibers in the thalamus are almost entirely limited to the central portion of the medial dorsal nucleus (B), with little to no FN-MdV fibers observed in other thalamic nuclei (C-F). G-H) In the midbrain, sparse fibers are observed in the mRT (G), with stronger expression observed in the LDTg (H). I-L) In the brainstem, extensive fibers are observed in regions including the PnC (I) and PCRT (J), as well as vestibular nuclei, including the SpVe (K). As was noted for FN-SC neurons, there is little evidence of FN-MdV neurons projecting to the spinal cord (L). MDC: Medial dorsal nucleus, central portion. All other abbreviations are as for Fig. 1. All images are taken from sagittal sections. Scale bars 50µm.

**Figure 6.**
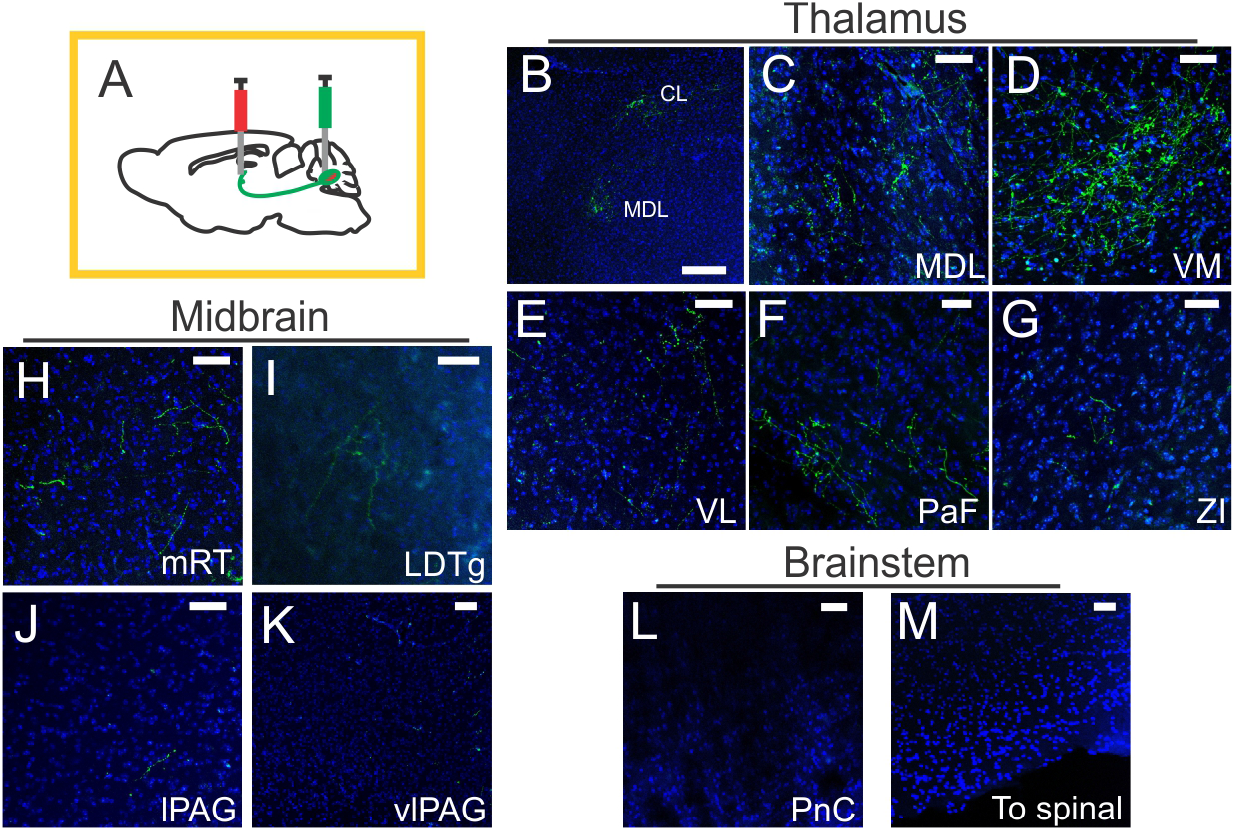
Collaterals from FN-CL neurons are observed in additional thalamic nuclei. A) Schematic of injection targeting for FN-CL neurons. B-G) In addition to the CL nucleus, FN-CL neurons also send projections to other thalamic nuclei, including MDL (C), and VM (D) thalamic nuclei. Sparse fibers are also observed in the PaF (E) and ZI (G). Note that panel B provides a lower magnification image of the thalamus, illustrating the patches of expression in the CL and MDL. H-K) Limited expression is observed in the midbrain. L) Essentially no fibers are observed in the brainstem, including PnC. L) As with FN-SC and FN-MdV neurons, there is no evidence of FN-CL neurons sending projections to the spinal cord. Abbreviations are as for Fig. 1. All images are taken from sagittal sections. Scale bars 50µm.

**Figure 7.**
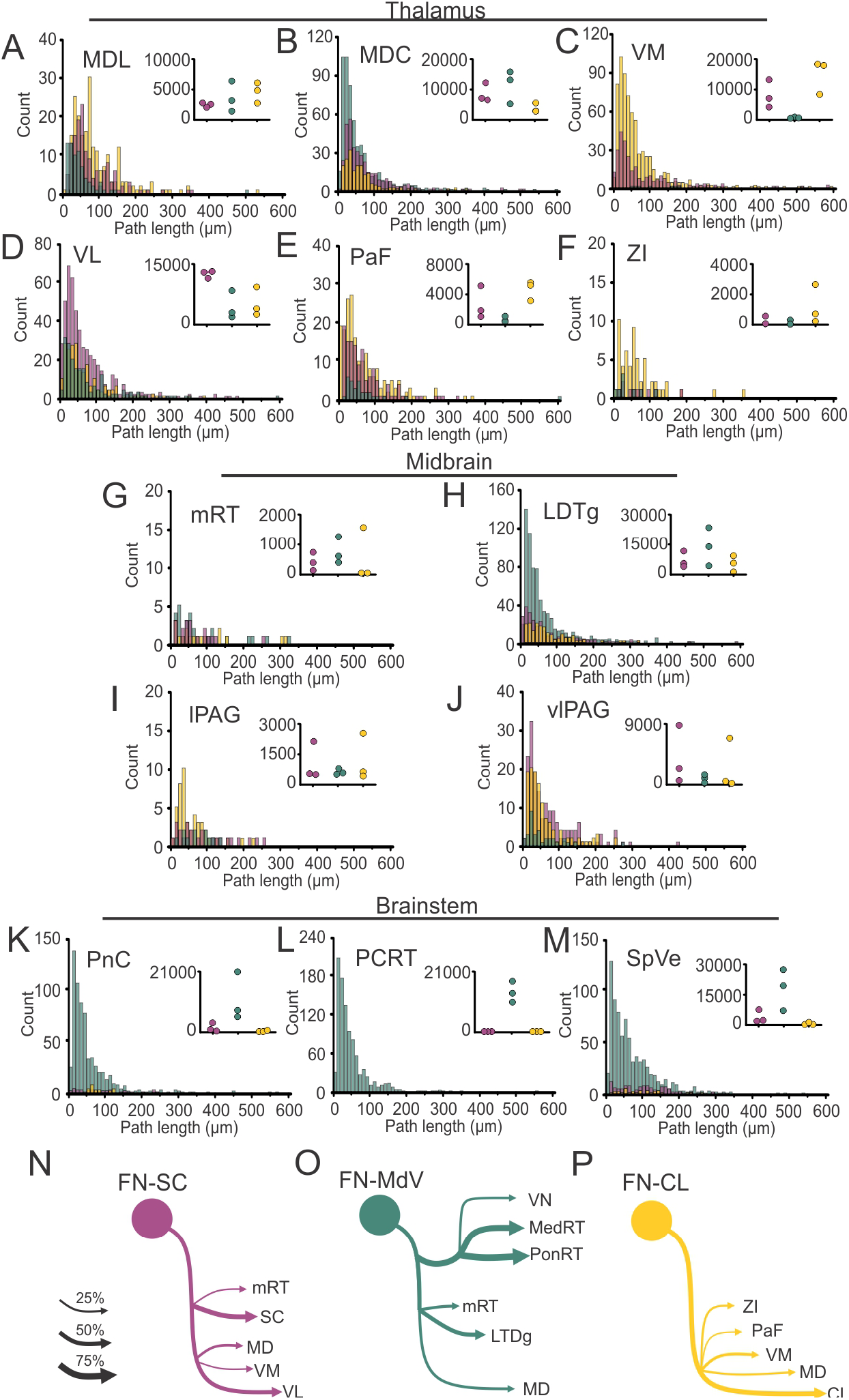
Fastigial outputs form largely segregated, but somewhat overlapping, channels. A-F) Histograms of path lengths of fibers in thalamic (A-F), midbrain (G-J), and brainstem nuclei (K-M) for FN-SC (magenta), FN-MdV (teal), and FN-CL (yellow) neurons (n = 3 animals per group, dot plot insets indicate total path length for each animal across all groups). Note differences in y-axis scales for different regions. N-P) Schematic illustrations of brain regions receiving relatively strong inputs from FN-SC (N), FN-MdV (O), and FN-CL (P) neurons, with line thickness roughly indicating relative strength, compared to max-observed fastigial inputs to that area. VN: Vestibular nuclei including SpVe, SuVE, and MVe; note that quantification was only done for SpVe. MedRT: Medullary reticular nuclei including Gi and MdD; note that quantification was only done for MdV. PonRT: Pontine reticular nuclei including PnC, PnO, PCRT, and PCRTa; note that quantification was only done for PnC and PCRT. All other abbreviations are as for Fig. 1-6.

Labeling of FN-SC neurons (Fig. 4A) also resulted in labeled fibers in several thalamic nuclei, including the ventral lateral nucleus, and, to a lesser extent, the medial dorsal and ventral medial nuclei (Fig. 4B-D). Some expression was also observed in the parafascicular nucleus in one animal (Fig. 4E), and little to no observed in the zona incerta (Fig. 4F). In other regions of the midbrain, expression was extremely limited (Fig. 4G-J). Similarly, essentially no fibers were observed in the pontine reticular formation (Fig. 4K) or vestibular nuclei, and we saw no evidence of FN-SC fibers travelling towards the spinal cord (Fig. 4L).

In contrast to FN-SC neurons, FN-MdV neurons (Fig. 5A) send relatively limited inputs to the thalamus (Fig. 5B-F), with only the central portion of the medial dorsal nucleus showing dense FN-MdV fibers (Fig. 5B). Within the midbrain (Fig. 5G-H), strong fiber labeling was seen in the laterodorsal tegmental nucleus (Fig. 5H). The highest density of FN-MdV fibers was observed in the brainstem, with extensive terminals in regions of the pontine (in addition to medullary) reticular formation (Fig. 5I-J) and vestibular nuclei (Fig. 5K). This widespread brainstem expression also included the gigantocellular reticular nucleus, the oral part of the pontine reticular nucleus (PnO), the medial vestibular nucleus and the superior vestibular nucleus. FN-MdV fibers in the brainstem largely appeared limited to local terminals, as we did not observe strong evidence of FN-MdV fibers traveling to the spinal cord (Fig. 5L).

Labeling of FN-CL neurons (Fig. 6A) also resulted in labeling of fibers in additional thalamic nuclei, including the medial dorsal (Fig. 6B-C) and ventral medial (Fig. 6D) thalamic nuclei, with more limited projections to the parafascicular nucleus (Fig. 6F). Finally, FN-CL neurons also appear to send very sparse projections to the zona incerta (Fig. 6G). As with FN-SC neurons, FN-CL collaterals to the midbrain (Fig. 6H-K) and brainstem (Fig. 6L-M) were very limited. Finally, similar to FN-SC and FN-MdV neurons, we did not observe evidence of FN-CL neurons sending projections to the spinal cord (Fig. 6M).

Quantification of fibers from FN-SC, FN-MdV, and FN-CL neurons further supports that these three populations send projections preferentially to largely segregated – but partially overlapping – thalamic, midbrain, and brainstem nuclei (Fig. 7). Overall, of these three output channels, FN-CL projecting neurons provided the strongest projections to the thalamus (Fig. 2O, Fig. 7A-F), with the notable exception of the VL (Fig. 7D) which received more input from FN-SC neurons, and the MDC (Fig. 7B), which received inputs from all three channels, but the heaviest from FN-MdV neurons. In the midbrain, inputs from all three populations were largely limited and somewhat overlapping (Fig. 7G-J), with FN-MdV inputs to the laterodorsal tegmental nucleus being the most notable. While some fibers were found in the periaqueductal gray areas (Fig. 7I,J), these projections were extremely limited compared to the expression observed in animals with broad fastigial targeting (Fig. 1F). Finally, fastigial inputs to the brainstem were massively dominated by FN-MdV neurons, which send a high density of fibers throughout the medullary (Fig. 2) and pontine reticular formation (Fig. 7K-L), as well as the vestibular nuclei (Fig. 7M). Note that only a subset of targeted regions are illustrated in the figures and quantified in Figure 7.

Together, these results indicate that there are distinct output channels from the fastigial nucleus, and that FN-SC, FN-MdV, and FN-CL neurons comprise distinct populations, with different, largely segregated, sets of target regions. FN-SC and FN-CL neurons each send projections to unique thalamic nuclei, with more limited expression in the midbrain and brainstem (Fig. 7N, P). Conversely, FN-MdV neurons send numerous projections to several nuclei within the brainstem (Fig. 7O), and have extremely restricted projections to the thalamus. With this knowledge in hand, we were able to test the impact of optogenetic manipulation of these populations of neurons on seizures.

### On-demand excitation of fastigial neurons projecting to the central lateral nucleus, but not SC or MdV, robustly attenuates hippocampal seizures

We utilized our dual viral strategy to implement on-demand optogenetic interventions in epileptic mice, in order to determine whether fastigial influence over seizure activity can be achieved via FN-SC, FN-MdV, or FN-CL neurons. Chronically epileptic animals were implanted with recording electrodes in the hippocampus, and the local field potential signal was analyzed in real-time to enable detection of spontaneous electrographic seizure activity and subsequent closed-loop intervention (Armstrong et al., 2013). Light was delivered to the fastigial nucleus in animals expressing channelrhodospin (ChR2) selectively in FN-SC, FN-MdV, or FN-CL neurons. Light was delivered for half of detected seizure events, in a random fashion, allowing each animal to serve as its own internal control.

Optogenetic excitation of FN-SC neurons (Fig. 8A) did not appear to have any effect on hippocampal seizures (Fig. 8B-D). Across the population, optogenetic excitation of FN-SC neurons failed to inhibit hippocampal seizures (Fig. 8D, p = 0.46, Kolmogorov-Smirnov test. Inset, 8 ± 6% average reduction in seizure duration) with no significant effect of light observed in any FN-SC animal tested (Fig. 8D inset, not significant in 6 of 6 animals). We also observed no effect on time to next seizure when modulating FN-SC neurons (p = 0.31, Wilcoxon signed rank test, n = 6 animals).

**Figure 8.**
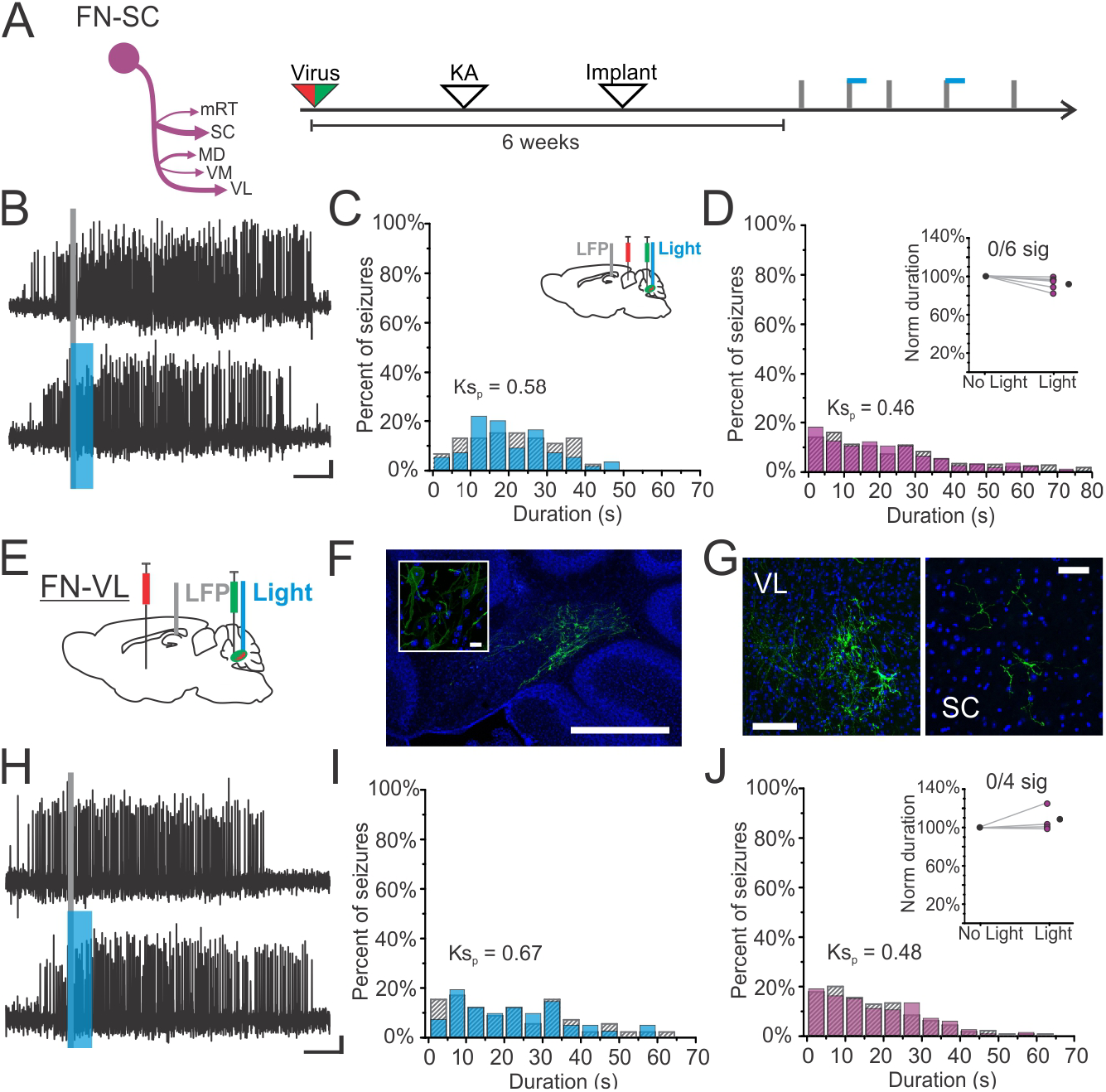
On-demand optogenetic excitation of FN-SC neurons fails to attenuate hippocampal seizures. A) FN-SC output channel and experimental design schematics. Mice received dual viral injections and underwent epilepsy induction, followed by implantation of electrodes in the hippocampus and optic fibers in the fastigial nucleus for targeting FN-SC. B) Example seizure events detected (gray line) on-line that were randomly elected to either not receive light (top trace) or receive on-demand light delivery (bottom trace) to FN-SC neurons. Blue bar indicates timing of light delivery. C) Histograms of post-detection seizure durations for events not receiving light (hashed bars) versus those receiving light (blue bars) in an example animal with targeting of FN-SC neurons, illustrating no significant effect of light delivery on hippocampal seizure duration (p = 0.58, two sample Kolmogorov-Smirnov test). Inset indicates experimental schematic of viral targeting, electrode recording, and optogenetic stimulation. D) Population histograms of post-detection seizure durations across FN-SC mice (n=600 events, from six animals), showing no significant effect of light delivery (p = 0.46, two sample Kolmogorov-Smirnov test). Inset: 0 out of 6 FN-SC mice (p > 0.05, two sample Kolmogorov-Smirnov test) show a significant effect of light delivery (black circle indicates mean). E) Additional targeting strategy, using retrograde viral injection into the VL nucleus of the thalamus. F) Dual viral targeting of FN-VL neurons labels neurons in the caudal portion of the fastigial nucleus (inset shows higher magnification image), as well as fastigial fibers in both the VL nucleus (G, left) and SC (G, right). H) Example seizure events detected (gray line) on-line that were randomly elected to either not receive light (top trace) or receive on-demand light delivery (bottom trace) to FN-VL neurons. I) Seizure durations from an example animal with targeting of FN-VL neurons, illustrating no significant effect of light delivery (p = 0.67, two sample Kolmogorov-Smirnov test). J) Population histograms of post-detection seizure durations across FN-VL mice (n=400 events, from four animals), showing no significant effect of light delivery (p = 0.48, two sample Kolmogorov-Smirnov test). Inset: 0 out of 4 FN-VL mice show a significant effect of light delivery. Scale bars: 5 sec, 0.05mV (B, H); 500µm (F, inset 10µm); 100µm (G, left); 50µm (G, right).

Notably, because we targeted the somata of FN-SC neurons, we modulated the entire output channel with this approach. This would suggest that these fastigial outputs to other areas targeted by FN-SC neurons (e.g., ventral lateral thalamus) are also unlikely to contribute meaningfully to the seizure suppressive effects of fastigial modulation. To further explore this possibility, we utilized the same dual viral approach to target fastigial neurons traveling to the ventral lateral nucleus of the thalamus (Fig. 8E-F), which project to both the ventral lateral nucleus (Fig. 8G, left) and the superior colliculus (Fig. 8G, right). As with FN-SC modulation, on-demand excitation of FN-VL neurons failed to attenuate hippocampal seizures (Fig. 8H-J), with no significant effect of light delivery observed at the population (Fig. 8J, p = 0.48, Kolmogorov-Smirnov test. Inset, average 8 ± 13% *increase* in seizure duration) or individual animal level (Fig. 8J inset, not significant in 4 out of 4 animals). Together, these results suggest that the FN-SC/FN-VL outputs do not mediate the seizure suppression observed with broadly targeting FN output neurons.

We next assessed whether excitation of FN-MdV neurons would have any effect on hippocampal seizures (Fig. 9A). Similar to FN-SC neurons, optogenetic excitation of FN-MdV neurons did not robustly attenuate hippocampal seizures (Fig. 9B-D). Across the population, a modest, albeit statistically significant effect of light was observed (Fig, 9D, p = 0.046, Kolmogorov-Smirnov test, Inset: average 5 ± 17% reduction in post-detection seizure duration). However, at the individual animal level, only 1 out of 6 animals showed a significant reduction in seizure duration (Fig. 9D inset arrow; Fig.9E). In this one animal, the measured reduction in seizure duration appeared to be more of a pause in seizure activity instead of a true attenuation (Fig 9F-H), indicating that optogenetic excitation of FN-MdV neurons is insufficient to truly attenuate hippocampal seizures. As with FN-SC neurons, these results also suggest that other areas targeted by FN-MdV neurons (e.g. vestibular nuclei, MDC, pontine reticular formation) are unlikely to be driving fastigial control of hippocampal seizures.

**Figure 9.**
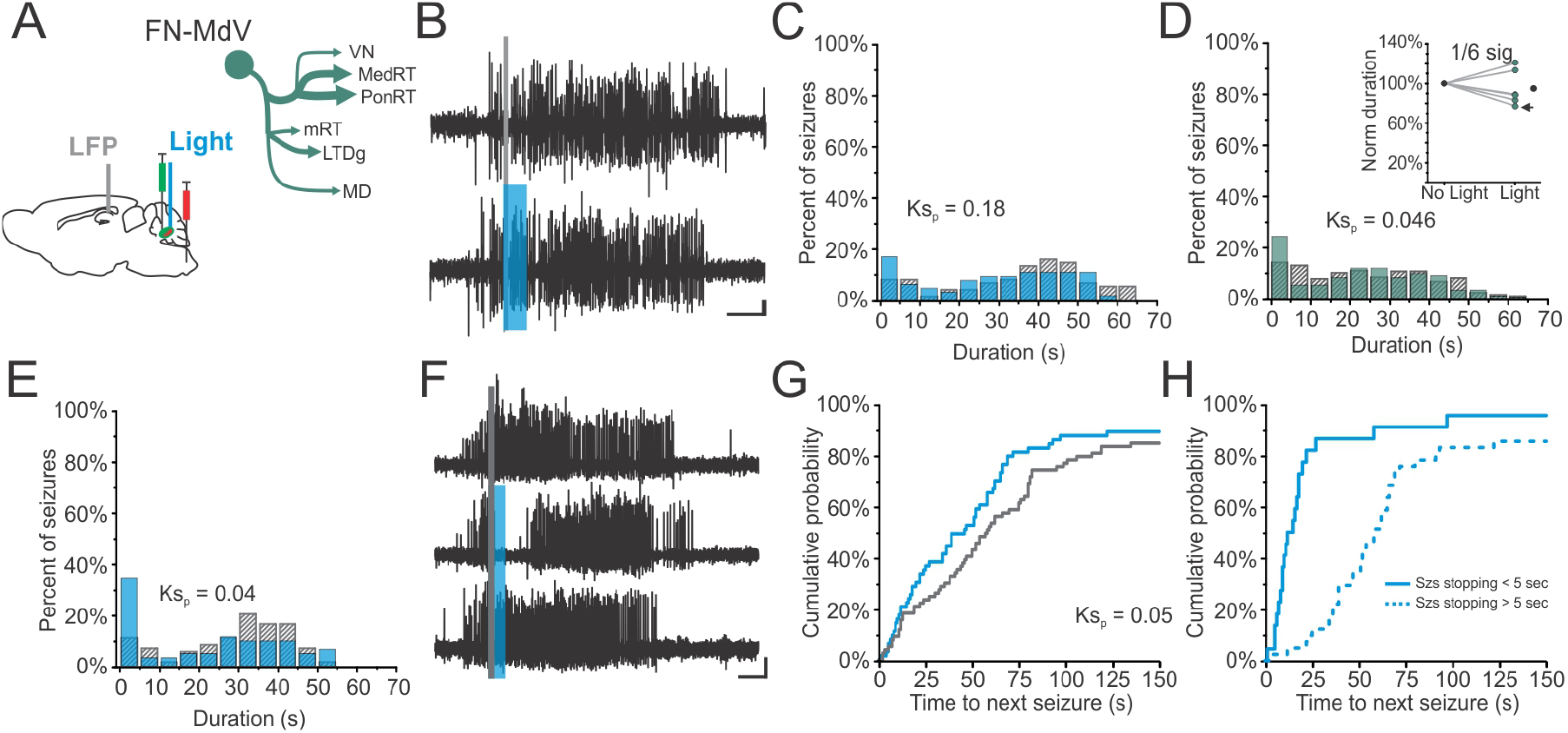
On-demand optogenetic excitation of FN-MdV neurons fails to attenuate hippocampal seizures. A) FN-MdV output channel and experimental schematic for targeting. B) Example seizure events detected (gray line) on-line that were randomly elected to either not receive light (top trace) or receive on-demand light delivery (bottom trace) to FN-MdV neurons. Blue bar indicates timing of light delivery. C) Seizure durations from an example animal with targeting of FN-MdV neurons, illustrating no significant effect of light delivery (p = 0.18, two sample Kolmogorov-Smirnov test). D) Population histograms of post-detection seizure durations across FN-MdV mice (n=600 events, from six animals), showing a marginally significant effect of light delivery (p = 0.046, two sample Kolmogorov-Smirnov test). Inset: 1 out of 6 FN-MdV mice show a significant effect of light delivery (indicated by arrow). E) Histograms of post-detection seizure durations for events not receiving light (hashed bars) versus those receiving light (blue bars) in the one FN-MdV animal showing a significant effect of light delivery (p = 0.04, two sample Kolmogorov-Smirnov test; 23% reduction in post-detection seizure duration with light in this animal). F) Example seizure events that were randomly elected to either not receive light (top trace) or receive light delivery (middle and bottom traces) to FN-MdV neurons in that one animal, illustrating that the effect of light delivery, when present, is more of a pause in seizures than a true attenuation. G) Cumulative probabilities of time to next seizure for seizure events receiving light (blue trace) and those not receiving light (gray trace), illustrating a trend towards a decrease in time to next seizure for events receiving light in this animal (p = 0.05, two sample Kolmogorov-Smirnov test; overall, a 13% reduction in time to next seizure). H) Seizure events which received light and stopped within 5 seconds tended to have a shorter time to next seizure (solid blue line) versus those that did not stop within 5 seconds (dotted blue line), again suggesting brief interruptions to seizures, rather than robust inhibition in this animal. Note that no light seizure events are not plotted, as only 12 no light seizures lasted less than 5 seconds. Scale bars: 5 sec, 0.05mV (B, F).

In contrast to FN-SC and FN-MdV targeting, optogenetic excitation of FN-CL neurons *was* sufficient to attenuate seizures (Fig. 10A-C). Across the population, targeting FN-CL neurons robustly and consistently attenuated hippocampal seizures (Fig. 10D, p < 0.001, Kolmogorov-Smirnov test) with all animals (7 of 7; Fig. 10D, Inset, average 34 ± 17% reduction in post-detection seizure duration) showing a significant effect of light delivery. This data indicates that on-demand activation of fastigial neurons projecting to the CL nucleus of the thalamus is sufficient for seizure inhibition. This finding is particularly striking given the lack of effect in either FN-SC or FN-MdV animals; note that FN-MdV animals not only had extensive fibers to numerous areas, but also had considerably more FN neurons labeled than FN-CL animals (Fig. 3), indicating that this effect cannot be explained by relative expression levels.

**Figure 10.**
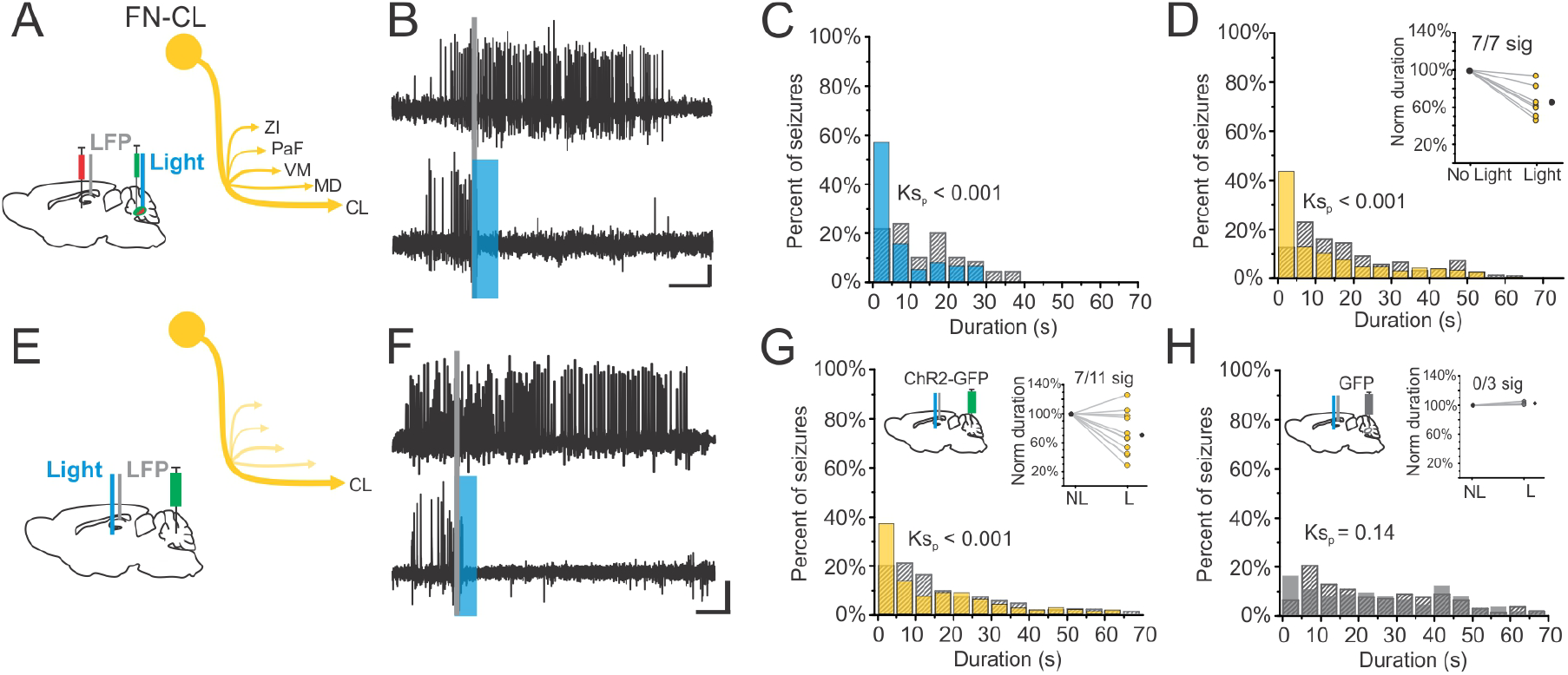
On-demand optogenetic activation of FN-CL neurons robustly attenuates hippocampal seizures. A) FN-CL output channel and experimental schematic for targeting. B) Example seizure events detected (gray line) on-line that were randomly elected to either not receive light (top trace) or receive on-demand light delivery (bottom trace) to FN-CL neurons. Blue bar indicates timing of light delivery. C) Seizure durations from an example animal with targeting of FN-CL neurons, showing a significant effect of light delivery (p < 0.001, two sample Kolmogorov-Smirnov test). D) Population histograms of post-detection seizure duration events across FN-CL mice (n=700 seizure events, from seven mice), showing a significant effect of light delivery (p < 0.001, two sample Kolmogorov-Smirnov test). Inset: Normalized seizure duration for light versus no light across FN-CL mice, with 7 out of 7 mice (100%) showing a significant reduction at the animal level. Black dot denotes average. E) To directly target FN fibers in the CL nucleus, mice were injected in the fastigial nucleus with virus encoding channelrhodopsin, followed by intrahippocampal kainic acid for induction of epilepsy, and then implanted with electrodes in the hippocampus and optic fibers targeting the central lateral nucleus for on-demand interventions. F) Example detected seizure events that were randomly elected to either not receive light (top trace) or receive light delivery (bottom trace) to fastigial fibers in the CL nucleus. Blue bar indicates timing of light delivery. G) Population histograms of post-detection seizure duration events across FN-CL mice (n=1100 seizure events, from eleven mice), showing a significant effect of light delivery (p < 0.001, two sample Kolmogorov-Smirnov test). Inset: Normalized seizure duration for light versus no light across CL fiber targeting mice, with 7 out of 11 mice showing a significant reduction at the animal level. Block dot indicates average. H) No effect of light delivery to the CL is observed in control animals (n = 300 seizure events from 3 mice, p = 0.14, two sample Kolmogorov-Smirnov test). Scale bars: 5 sec, 0.05mV (B, F).

In separate animals, we directly targeted the terminals of fastigial neurons in the CL (Fig. 10E-F). Despite caveats mentioned above associated with targeting terminals, we found significant inhibition of seizures with this approach (Fig. 10G, average 27 ± 9% reduction, p < 0.001 Kolmogorov-Smirnov test; inset: significant at the individual animal level in 7 of 11 animals). This further supports that activation of FN-CL neurons is sufficient to attenuate hippocampal seizures.

Confirming that the effect of light delivery to the CL was mediated via opsin-activation, no effect was observed for opsin negative animals (injected with a virus for expression of GFP only) receiving light to the CL (Fig. 10H).

Across the populations, in contrast to effects seen in an FN-MdV animal, there was no effect on time to next seizure when targeting FN-CL neurons (p = 0.297, Wilcoxon signed rank test), nor when targeting fastigial terminals in the CL (p = 0.240, Wilcoxon signed rank test). At the individual animal level, only one animal showed a significant effect of light delivery on time to next seizure, but it was an *increase* in time to next seizure with light (rather than a decrease). This suggests that FN-CL neurons produce a true seizure cessation (rather than brief pause).

Together, these results show that fastigial attenuation of hippocampal seizures can be achieved by selectively exciting FN-CL neurons, but not by exciting FN-SC/FN-VL or FN-MdV neurons. Our findings argue against the need for broad modulation of fastigial outputs, as activation of fastigial outputs to the CL thalamus, selectively, is sufficient to inhibit seizures. More broadly, our results highlight the concept of distinct fastigial output channels (Fig. 7N-P), and illustrate that the existence of such distinct output channels can have important functional consequences (Fig. 8-10).

## DISCUSSION

Using viral approaches coupled with neuroanatomical tracing and on-demand optogenetics, this study reveals that a specific fastigial output channel is sufficient for attenuation of hippocampal seizures. While the fastigial nucleus projects to a great many different target regions, we found that outputs are organized into segregated channels. Specifically, fastigial neurons which project to the SC also project heavily to the ventral lateral thalamus. Fastigial neurons projecting to the MdV also target central regions of the medial dorsal thalamic nucleus, and send dense, widespread, projections to brainstem reticular and vestibular nuclei. Finally, fastigial neurons that project to the CL thalamus also project to the ventral medial and medial dorsal thalamic nuclei, and to a lesser extent areas including zona incerta and the parafascicular nucleus. We found that excitation of FN-CL neurons is sufficient to attenuate seizures, whereas excitation of fastigial neurons projecting to the SC or MdV has no effect on seizures. Together, these results indicate that fastigial control of hippocampal seizures does not require broad, concurrent manipulation of several fastigial output channels; excitation of the fastigial output channel to CL thalamus is sufficient. Notably, these findings also illustrate the functional relevance of distinct fastigial output channels, and the potential usefulness of understanding cerebellar circuitry at this level.

We previously demonstrated that fastigial excitation can robustly attenuate hippocampal seizures (Streng and Krook-Magnuson, 2020b). However, the lack of direct projections to the hippocampus (Strick et al., 2009; Rochefort et al., 2013; Bohne et al., 2019; Watson et al., 2019; Krook-Magnuson, 2020) requires that seizure cessation be mediated via connections with at least one intermediate region. Despite being densely connected with a very large number of downstream targets (>60 total as estimated by Fujita *et al* 2020), we demonstrate that the fastigial output channel to the CL nucleus of the thalamus is sufficient for seizure suppression. An intralaminar nucleus considered to be part of the higher order thalamus (Saalmann, 2014), the CL nucleus has been implicated in attention (Schiff et al., 2013), working memory (Wyder et al., 2004), and arousal (Van der Werf et al., 2002). Especially relevant in the context of epilepsy and seizures, it is hypothesized that the intralaminar thalamic nuclei serve to synchronize -- and desynchronize -- the cerebral cortex in order to coordinate activity of cortical neurons and networks (Saalmann, 2014). Focal limbic seizures modulate neuronal firing in the CL nucleus, causing both an overall decrease in their activity levels and an increase in bursting (Feng et al., 2017). Additionally, electrical stimulation of the CL nucleus during acutely evoked hippocampal seizures can induce cortical desynchronization and improve measures of consciousness and arousal (Gummadavelli et al., 2015; Kundishora et al., 2017; Xu et al., 2020), suggesting this nucleus can play an active role in both participating in and influencing seizure networks. Our data illustrate that perturbing activity in the CL nucleus may be able to inhibit seizures, as activation of FN-CL neurons attenuated spontaneous hippocampal seizures (Fig. 10). Notably, we recorded seizures in the hippocampus near the previous site of kainic acid injection and the presumed seizure focus, likely indicating that the CL is ultimately able to impact “upstream” hippocampal seizure activity. This may be through broad impacts on cortical synchrony (as noted above), or specific effects e.g. on the anterior cingulate cortex (Van der Werf et al., 2002), which is reported to have direct projections to the hippocampus (Rajasethupathy et al., 2015).

While our work utilized the intrahippocampal kainate mouse model of temporal lobe epilepsy, the ability of the cerebellum to influence seizures may extend to those arising in regions outside the hippocampus (reviewed in (Streng and Krook-Magnuson, 2020a)), including thalamocortical absence seizures (Kros et al., 2015). It remains unknown whether cerebellar influence over different seizure networks is also mediated by neurons projecting to the CL. In a recent study examining thalamocortical absence seizures, activation of cerebellar nuclei terminals in the thalamus was able to attenuate spike-and-wave discharges, but little appeared to be mediated via the CL nucleus in particular (Eelkman Rooda et al., 2021). Indeed, that work tentatively concluded that cerebellar inhibition of absence seizures was mediated via multiple nuclear output (channels) working together. This is in stark contrast to our findings; differences in outcomes may be due to epilepsy type, cerebellar nucleus targeted, or experimental methods (i.e., we primarily targeted somata within the cerebellar nucleus to avoid difficulties with selective and sufficient terminal activation).

It is important to note that our experiments examine the FN-CL output channel, which has collaterals to additional areas. Even for our fiber targeting experiments, in which light was delivered directly to the CL, we cannot rule out potential antidromic activation. As such, other areas FN-CL neurons project to, including the VM nucleus of the thalamus, may contribute to the anti-seizure effects shown here. The VM projects to cortical areas including anterolateral motor cortex (Guo et al., 2018) and infralimbic prefrontal cortex (Sieveritz and Arbuthnott, 2020), and can target inhibitory neurons (and in particular, layer 1 neurogliaform cells) (Armstrong et al., 2012; Overstreet-Wadiche and McBain, 2015; Anastasiades et al., 2021). Given the potential interest of basal ganglia circuitry to seizures (Iadarola and Gale, 1982; Garant and Gale, 1987; Wicker et al., 2019), it is also of interest that the VM receives inhibitory input from pars reticulata (Kase et al., 2015). Additionally, neurons in this nucleus are highly modulated by spike-and-wave discharges during absence seizures (Paz et al., 2007), and targeting cerebellar terminals in the VM can provide some inhibition of spike-and-wave absence seizures (Eelkman Rooda et al., 2021). However, the VM is also targeted by FN-SC neurons (Fig. 7N), albeit to a lesser extent, and activation of FN-SC neurons did not inhibit hippocampal seizures (Fig. 8). Similarly, while the MD is an additional target of FN-CL neurons and is of potential interest to seizure control (Cassidy and Gale, 1998; Bertram et al., 2008; Sloan et al., 2011), the MD also receives input from FN-SC and FN-MdV, which were ineffective in inhibiting seizures. Therefore, either the VM and MD are playing little to no role in the seizure-inhibition effects of FN-CL neurons, FN-SC/FN-MdV projections to the VM/MD are insufficient to achieve seizure inhibition, and/or FN-SC/FN-MdV neurons and FN-CL neurons project to different populations of VM/MD neurons.

It is important to note that our results firmly show that not all thalamic nuclei which receive fastigial input provide seizure inhibition effects. Specifically, we show that activation of FN-SC neurons, which have collaterals to the VL thalamus, does not inhibit seizures. We additionally directly targeted FN-VL neurons, and again saw no inhibition of seizures. Therefore, fastigial inhibition of seizures is not mediated by fastigial-thalamic projections in a generic sense, but rather by FN-CL neurons specifically.

In this regard, in addition to identifying key fastigial outputs necessary for seizure cessation, our results provide important evidence as to which targets may *not* be involved in mediating the anti-seizure effects of direct fastigial modulation. Given that direct optogenetic modulation of the SC is highly effective in disrupting seizures in several rodent models of different types of epilepsy (Soper et al., 2016), we initially reasoned that fastigial projections to the SC might be mediating fastigial inhibition of seizures. However, somewhat surprisingly, on-demand optogenetic excitation of FN-SC neurons failed to inhibit hippocampal seizures. Previous work has examined SC influence on seizures not arising in the hippocampus *per se* (Iadarola and Gale, 1982; Gale et al., 1993; Soper et al., 2016). As such, one possibility is that the SC lacks the ability to influence hippocampal seizures specifically. Alternatively, as with FN-SC projections to the VL or MD thalamus, FN projections to the SC may target the wrong population of SC neurons (e.g., ones that do not mediate seizure suppression), or may simply provide insufficiently strong input to the SC. Regardless, fastigial control of hippocampal seizures does not appear to be mediated via the SC.

We also show that excitation of fastigial projections to the medullary reticular formation fails to attenuate seizures. This argues against the hypothesis that seizure suppression is mediated by a reticular-formation-induced brain state change (Pfaff et al., 2012; Ewell et al., 2015; Khan et al., 2018; Purnell et al., 2018; Sakai, 2018; Streng and Krook-Magnuson, 2020a). Given the extensive fibers observed with FN-MdV labeling, as well as the distribution of cell bodies throughout the fastigial nucleus, the FN-MdV output channel may actually be comprised of multiple sub-channels. These sub-channels may contain different aggregates of additional downstream targets, such as those described previously (Teune et al., 2000). None appear to mediate seizure inhibition. Our results additionally indicate that other regions receiving strong input from FN-MdV neurons, including the central portion of the MD thalamus or the vestibular nuclei, are unlikely to be significantly contributing to suppression of hippocampal seizures.

While we cannot rule out that other FN output channels not tested in this study may also be able to inhibit seizures, the robust effect of FN-CL modulation, coupled with the lack of an effect when modulating either FN-SC or FN-MdV output channels, strongly argues against suppression of hippocampal seizures requiring coordinated changes across a large number of fastigial targets. Rather, there are particular pathway(s) – starting with the FN-CL output pathway -- by which the cerebellum influences the hippocampus and hippocampal seizures. While a different pathway and/or output channel may be at play in healthy animals, it is worth noting that temporally precise modulation of hippocampal activity is observed with stimulation of cerebellar neurons in non-epileptic animals (Choe et al., 2018; Zeidler et al., 2020). Additionally, multisynaptic rabies tracing, following injections into the hippocampal dentate gyrus (again, in non-epileptic animals), results in labeling of neurons in the fastigial nucleus (Watson et al., 2019), futher supporting a (multisynaptic) connection between these structures. Notably, this rabies labeling is seen in the *caudal* portion of the fastigial nucleus -- the same region in which we observe FN-CL neurons (Fig. 2Ciii), suggesting that hippocampal activity is influenced by this subpopulation of fastigial neurons.

The fastigial nucleus, considered part of the functional division of the cerebellum known as the spinocerebellum, has canonically been considered to be mostly involved in motor functions of proximal muscles and eye movements (Purves et al., 2001). While early work (Angaut and Bowsher, 1970; Batton et al., 1977; Bentivoglio and Kuypers, 1982; Angaut et al., 1985) has characterized some of the projection targets of cerebellar nuclei, including several thalamic and brainstem nuclei, and suggested some segregation of outputs based on somatic location within the fastigial nucleus (Bentivoglio and Kuypers, 1982; Noda et al., 1990; Fuchs et al., 1993; Teune et al., 2000), we are just beginning to appreciate the full diversity and complexity of fastigial projections to both motor and non-motor structures. A recent study examined output from fastigial neurons by subregion within the fastigial nucleus, and suggested 5 distinct output channels (Fujita et al., 2020), 3 of which bear many similarities with the populations we characterize here. In that study by Fujita and colleagues, one population of neurons in the caudal fastigial nucleus (and to some extent the caudal dorsolateral protuberance) projected to areas including the CL, MD, and VM thalamus (similar to our FN-CL channel). A second population of neurons in the caudal dorsolateral protuberance (and to some extent caudal fastigial) projected to areas including the VL and PaF thalamus, zona incerta, and SC (somewhat in keeping with our FN-SC channel). A third population of neurons in more rostral portions of the fastigial projected to areas including the medullary reticular formation, vestibular nuclei, and spinal cord (apparently encompassing some of our FN-MdV channel and additional channel(s)). In total, these experiments provide a wealth of insights into fastigial outputs. However, the experimental approaches taken did not allow segregation of output channels beyond immunohistochemical profile or general location within the nucleus – for example, it was not clear from those results if the same neuron projects to a single downstream area, to multiple downstream areas, or to all downstream areas identified as receiving input from a given region of the fastigial nucleus. Therefore, while Fujita *et al* identified a region of fastigial neurons which contains projections to the CL, ventral medial, and medial dorsal nuclei, we specifically show that fastigial neurons that project to the CL have collaterals to the ventral medial and medial dorsal thalamus – that is, the *same* neuron projects to multiple regions. Given the partial blurring of anatomical regions within the fastigial seen by Fujita and colleagues, it is also difficult to know from those studies if the output channels identified were non-overlapping. Indeed, our work suggests that while the FN-CL, FN-SC, and FN-MdV populations are distinct, there are areas of overlap which receive input from multiple output channels (e.g. the medial dorsal nucleus). We also note some additional slight differences in the inferred organization of outputs. Our results thus illustrate a high level of nuance in fastigial outputs, with an increased number of output channels than the five which were recently suggested, and where specific fastigial output channels send collateral projections to both convergent and divergent downstream targets.

The existence of individual fastigial neurons that project simultaneously to multiple, distinct, downstream regions is intriguing, as is their functional significance pertaining to ongoing, healthy, cerebellar function. One potential framework for understanding these simultaneous projections is the forward-internal-model hypothesis of cerebellar function, in which the cerebellum generates predictions about the sensory consequences of actions (Wolpert et al., 1998; Kawato, 1999; Popa and Ebner, 2018). These predictions of sensory consequences are used both to influence and correct ongoing processes as well as optimize future actions (for review, see (Popa et al., 2016)); fastigial collaterals could convey predictions to regions that would update ongoing motor commands and the motor controller. The existence of *segregated* channels may also be compatible with a multiple forward internal model framework; the cerebellum may implement forward internal models in motor and non-motor domains (Popa et al., 2014; Diedrichsen et al., 2019), via separate output channels. Alternatively, different regions and modules of the cerebellum may implement distinct functional roles (Diedrichsen et al., 2019). In either framework, FN-CL neurons may support more cognitive functions.

Together, our results shed important light on the necessary outputs for fastigial control of hippocampal seizures and demonstrate the complexity of fastigial outputs and projections. Rather than requiring broad modulation of fastigial outputs, we show that seizure suppression is achievable via FN-CL neurons, outlining new potential avenues for selective therapeutic interventions with minimized side effects. Our results also underscore the sophistication of the fastigial nucleus, whereby specific fastigial output channels, which project to specific downstream targets, carry distinct functional outcomes.

## Acknowledgments

Dr. Michael Benneyworth and the University of Minnesota Mouse Behavioral Core and Dr. Erin Larson and the University of Minnesota Optogenetics Core provided technical assistance and equipment. Laurel Schuck, Susan Khanpour, Madeline Marker, Jacob Weiner, Thai Loyd, Jack Redepenning, Isaac Hoff, Jane Yap, and Kayla Togneri of the Krook-Magnuson lab provided technical assistance.

## Funding

This work was supported in part by The Winston and Maxine Wallin Neuroscience Discovery Fund Award, an American Epilepsy Society Postdoctoral Fellowship (MLS), NIH R01-NS112518, a University of Minnesota McKnight Land-Grant Professorship award, and the University of Minnesota’s MnDRIVE (Minnesota’s Discovery, Research and Innovation Economy) initiative.

## Notes

The authors declare no competing financial interests.

### Competing Interest Statement

The authors have declared no competing interest.

